## supplemental figures for "Id2 GABAergic interneurons: a neglected fourth major group of cortical inhibitory cells"

**Fig. S1: scRNAseq gene heat maps (adapted from Yao et al., 2021)**
https://portal.brain-map.org/atlases-and-data/rnaseq

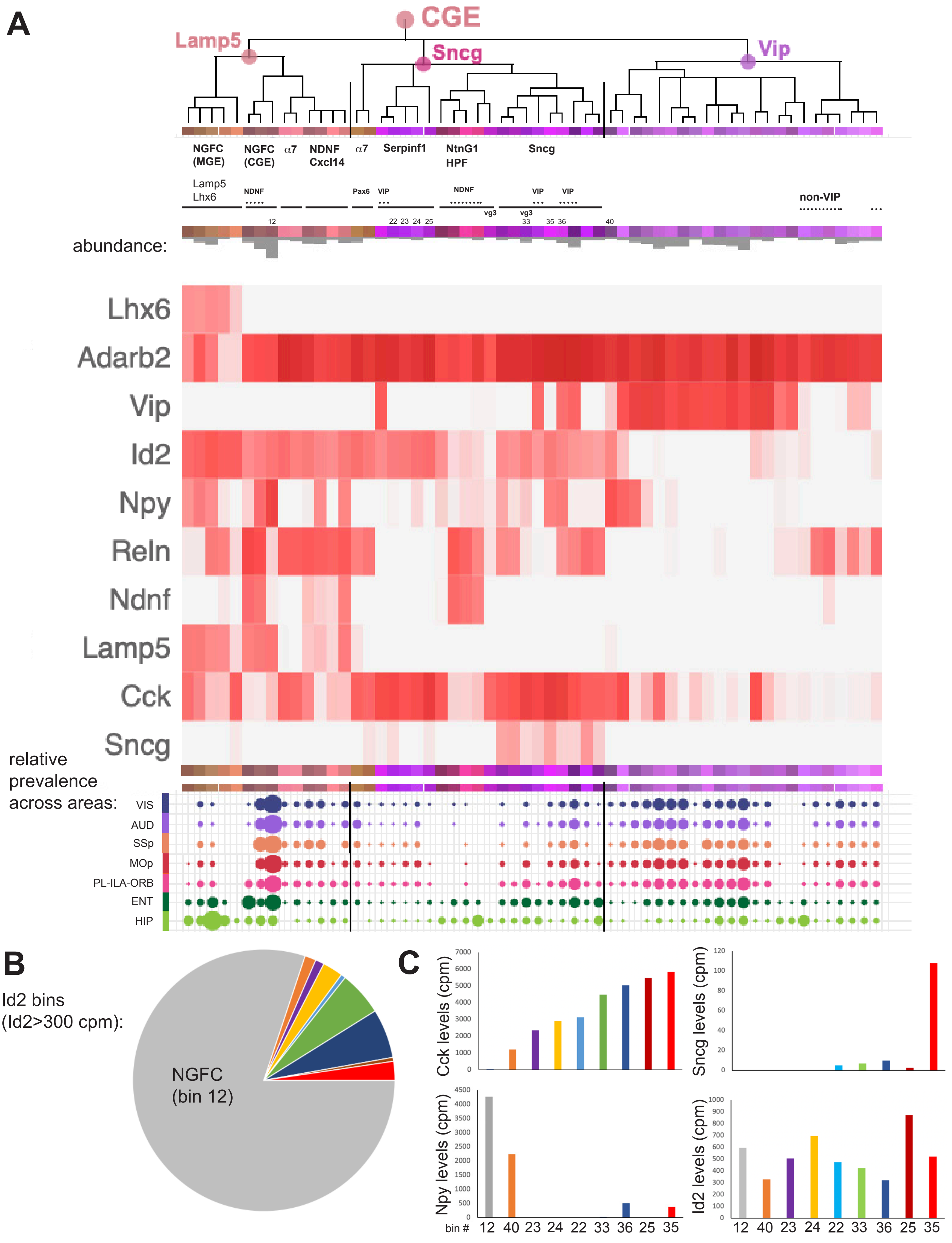

**Figure S1: CGE IN scRNAseq gene heat maps.** (**A**) Annotated dendrogram and gene heat maps based on IN scRNAseq data published by the Allen Institute (Yao et al. 2021; portal.brain-map.org/atlases-and-data/rnaseq). Heat maps illustrate mRNA expression levels (trimmed mean (25%-75%) LOG2(CPM+1)) for each gene listed: *Lhx6* (pan-MGE), *Adarb2* (pan-CGE), *Vip*, *Id2*, *Npy*, *Reln*, *Ndnf*, *Lamp5*, *Cck*, and *Sncg*. Based on our analysis of gene expression patterns, we added annotations beneath the major branches of the dendrogram to indicate the putative IN subtypes and markers that correspond with each set of color-coded bins. An abridged version of the schematic illustrating the relative prevalence of each IN bin type across different cortical areas and hippocampus from Yao et al., 2021 is aligned beneath the gene heat maps. (**B**) Pie chart of the nine IN bins where trimmed mean Id2 mRNA levels are >300 counts per million illustrates the main populations expected to be labeled in the Id2-CreER; Dlx5/6-Flpe; Ai65 intersectional cross. The number of cells in each bin was obtained from the Allen scRNAseq data portal (portal.brain-map.org/altlases-and-data/rnaseq) and includes cells isolated from throughout the cortex and hippocampus. (**C**) Plots of expression levels (counts per million) of *Cck*, *Sncg*, *Npy,* and *Id2* transcripts across the nine Id2 IN bins shown in (B). *Cck* exhibits variable levels of expression across the non-NGFC bins, with the highest levels in bin 35. *Sncg* is weakly expressed or absent in all but bin 35. Bin 12 (NGFC) shows the highest *Npy* level, as expected.

### Fig. S2: Prevalence and distribution of Id2 INs in prelimbic cortex

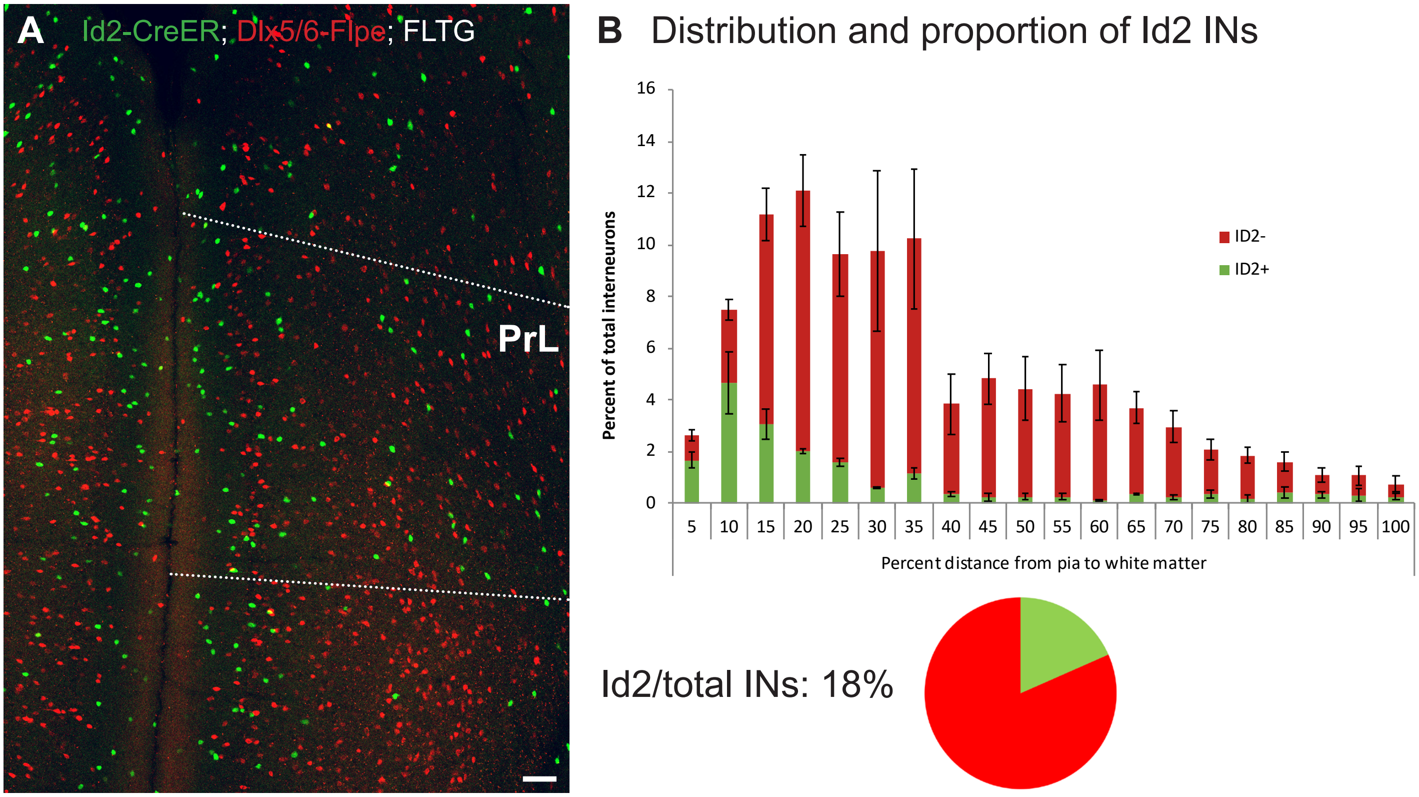

**Figure S2: Prevalence and distribution of Id2 INs in prelimbic cortex.** (**A**) Representative image of dual color labeling of Id2 INs (green) and non-Id2 INs (red) in the mPFC of the Id2-CreER; Dlx5/6-Flpe; FLTG cross. The prelimbic area (PrL) is denoted by the white dotted lines. The scale bar indicates 100 μm. (**B**) Quantification of the distribution and proportion of Id2 INs in PrL. Due to the difficulty in identifying precise layer boundaries in PrL, green and red cells were counted across 20 equally spaced bins across the pia to the white matter (438/2424 cells counted; n=3 brains; error bars indicate SEM). The overall fraction of Id2 INs/total INs was 18% (represented as a pie chart).

**Fig. S3:**
**Id2 INs show little overlap with Nkx2.1-lineage or VIP cells**

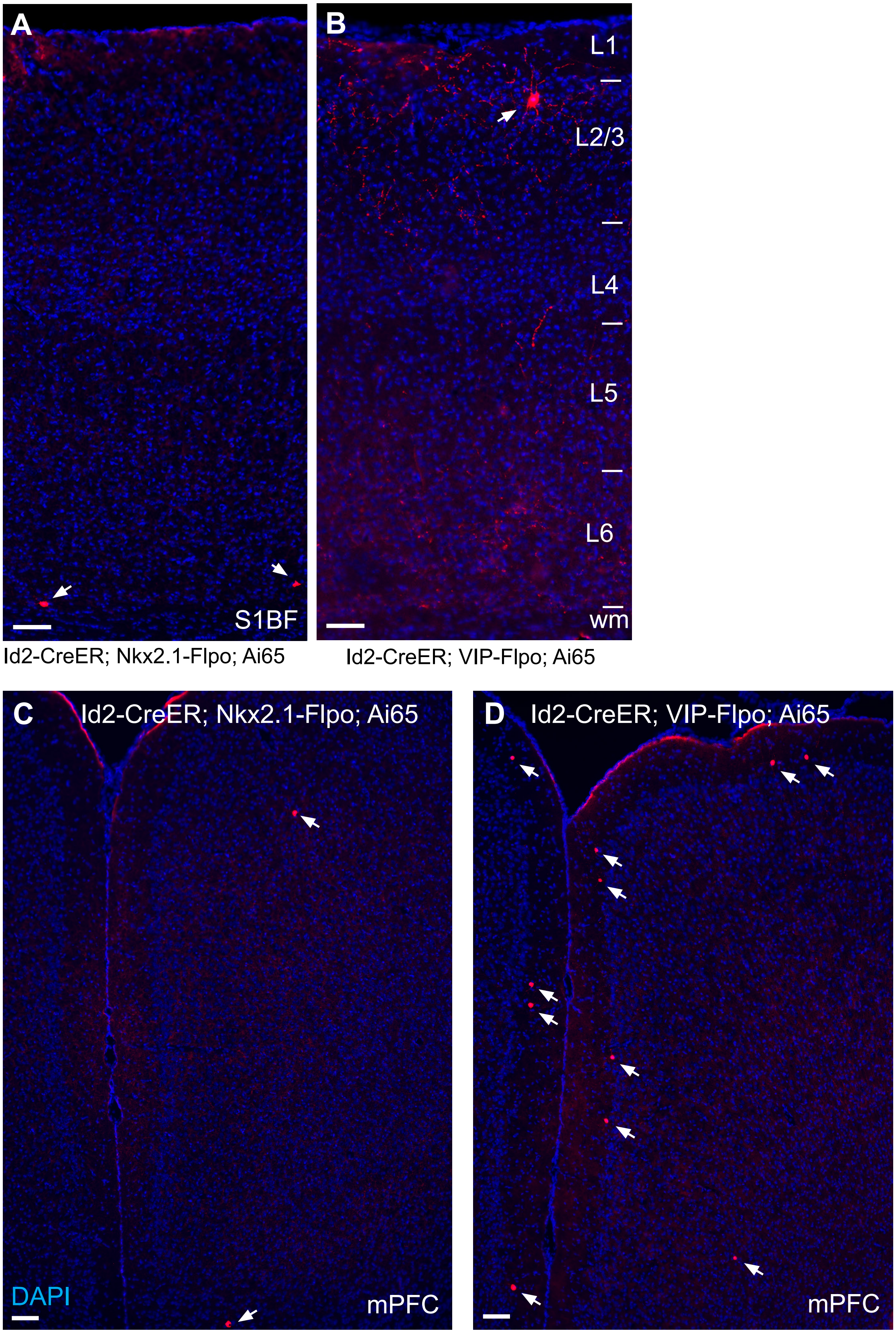

**Figure S3: Id2 INs show little overlap with Nkx2.1-lineage or VIP cells.** (**A**) Tissue section of the cortical column in S1BF of an Id2-CreER; Nkx2.1-Flpo; Ai65 brain. Arrows point to the two labeled cells in this field, both located in L6. (**B**) Tissue section of the cortical column in S1BF of an Id2-CreER; VIP-Flpo; Ai65 brain. The white arrow indicates the one labeled cell in this field, located in upper L2/3. Cell labeling was extremely sparse in both crosses. (**C**) Tissue section of the mPFC of an Id2-CreER; Nkx2.1-Flpo; Ai65 brain. Arrows point to the two labeled cells in this field. (**D**) Tissue section of the mPFC of an Id2-CreER; VIP-Flpo; Ai65 brain. The white arrows indicate the labeled cells in this field, mostly located along the L1/L2 border. DAPI counterstain (blue) is present in all images. All tissue sections are 20 $\mu$m thick. Scale bars indicate 100 $\mu$m.

#### Fig. S4: Id2 INs and NPY expression in S1BF, V1, and A1

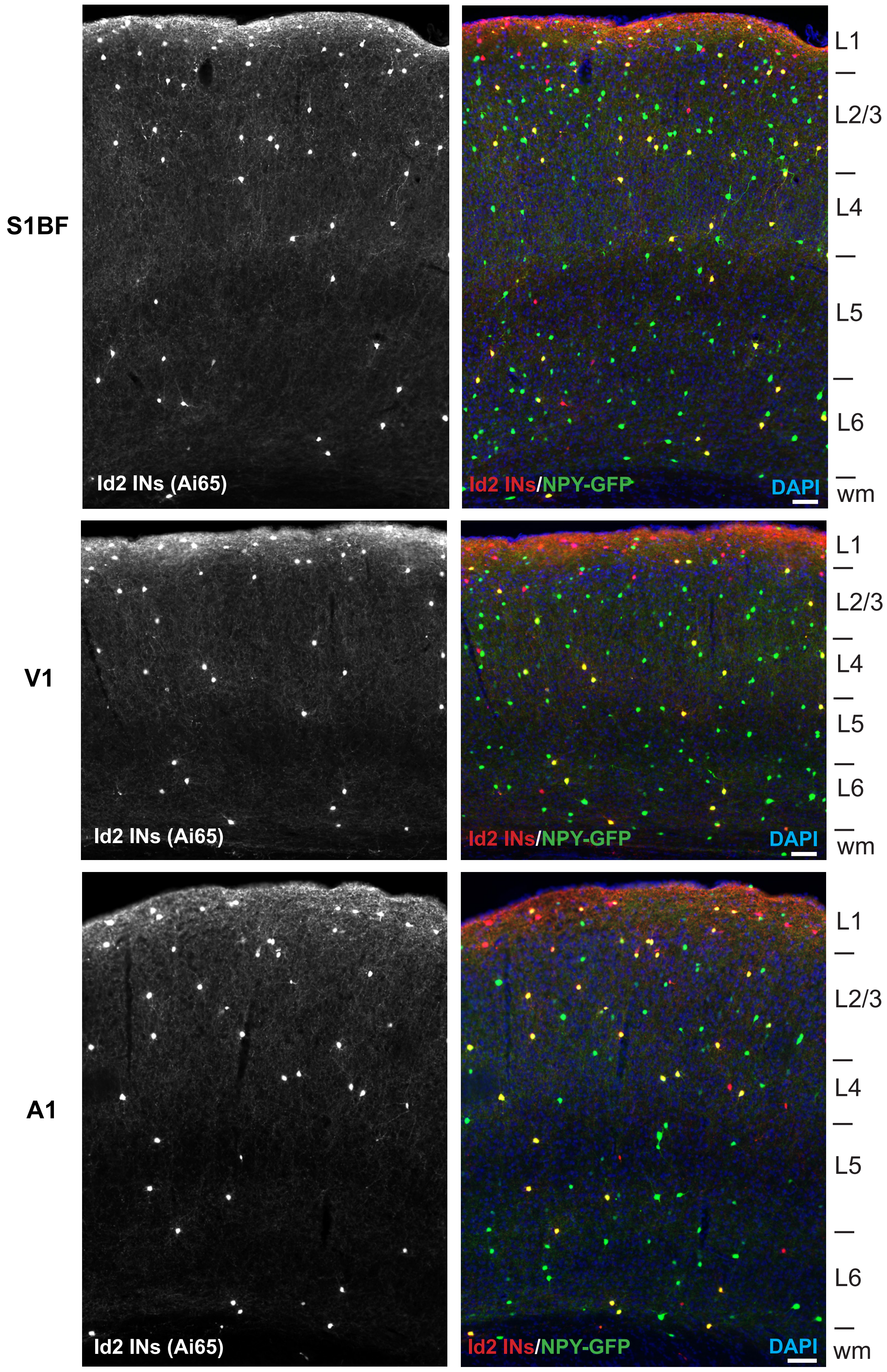

**Figure S4: Id2 INs and NPY expression in S1BF, V1, and A1.** Examples of Id2 IN labeling and NPY expression (tdTomato and NPY-hrGFP expression in Id2-CreER; Dlx5/6-Flpe; Ai65; NPY-hrGFP animals) in brain tissue sections from S1BF, V1 and A1. Id2 INs labeled with tdTomato are shown on the left panels, with the NPY-hrGFP signal included in the right panels from each cortical area. Red cells indicate Id2 INs without NPY expression, and yellow cells indicate Id2/NPY INs. The approximate locations of layer boundaries are labeled on the right. DAPI counterstain (blue) is present in all images. All tissue sections are 20 μm thick. Scale bars indicate 100 μm.

#### Fig. S5: Additional morphologies of Id2 INs

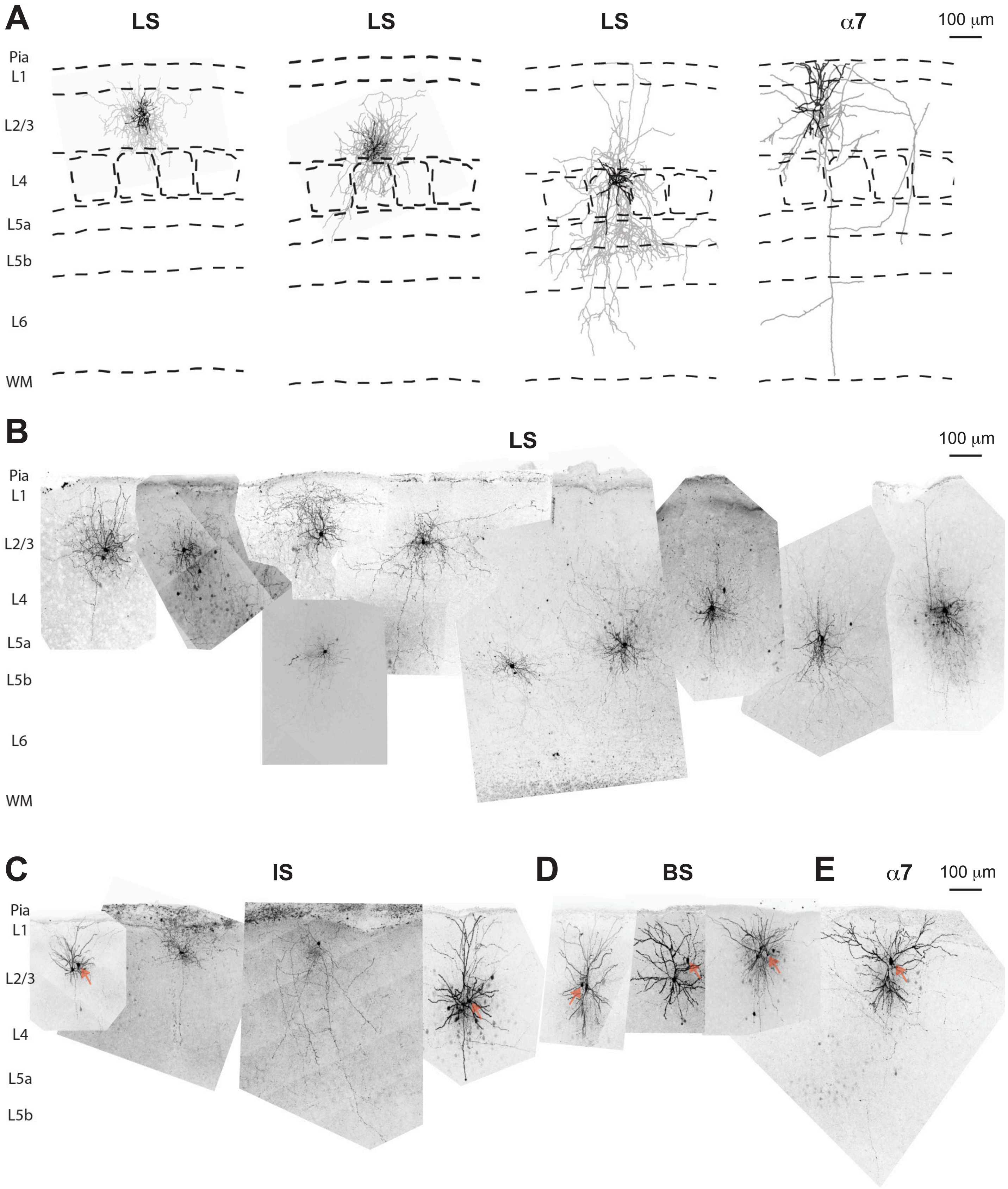

**Figure S5: Additional morphologies of Id2 INs.** (**A**) Reconstructions of three additional Id2 late-spiking (LS) INs and one Id2 alpha-7 ($\alpha$7) IN in L2-4 of S1BF. Approximate layer boundaries and barrel locations are indicated by dashed lines. (**B**) Images of ten filled Id2 LS INs aligned to their respective laminar locations. (**C**) Images of four filled Id2 irregular spiking (IS) INs and pyramidal cell recorded pairs located in L2/3. (**D**) Images of three filled Id2 burst spiking (BS) INs and pyramidal cell recorded pairs located in L2/3. (**E**) Image of an alpha-7 ($\alpha$7) IN and pyramidal cell recorded pair. Arrows in (**C**-**E**) indicate cell soma of the respective filled Id2 cells. Note: neuronal images are better appreciated by zooming in (especially axonal arbors).

**Fig. S6: Methodological considerations for assessing gene expression: The cautionary tale of CCK**

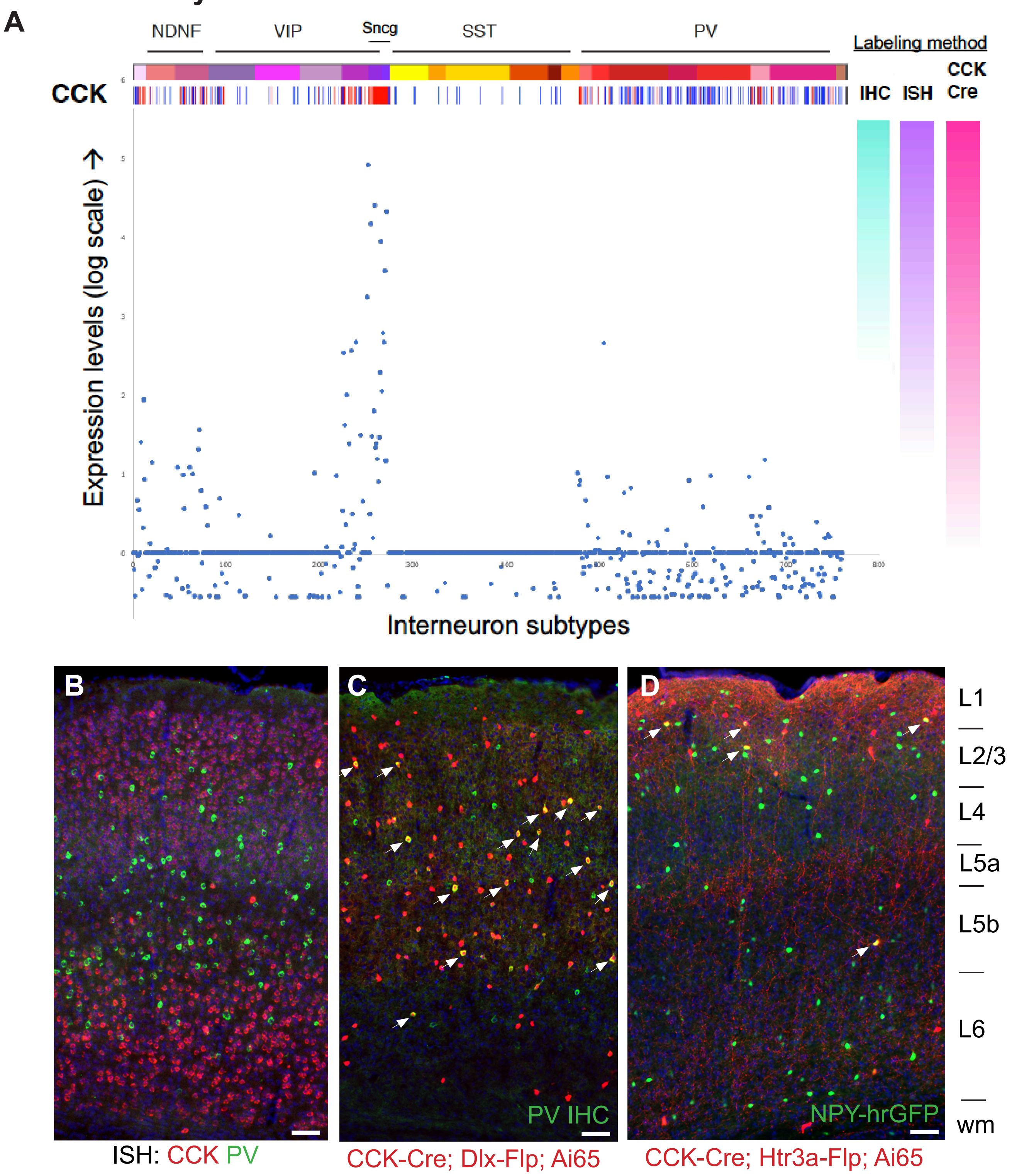

**Figure S6: Methodological considerations for assessing gene expression: the cautionary tale of CCK.** (**A**) Expression levels of *Cck* mRNA across all GABAergic INs from Tasic et al., 2016 (~3000 INs individually purified from V1), represented on a log scale. The top bar shows the IN subtype categories described in Tasic et al., 2016, with a heat map for *Cck* mRNA shown underneath. On the right, a schematic depiction of the hypothetical extent of cell labeling using immunohistochemistry (IHC), *in situ* hybridization (ISH), or CCK-Cre cumulative reporter labeling (e.g., Cck-Cre; Ai14) is shown. (**B**) Dual fluorescence *in situ* hybridization (FISH) for *Cck* (red) and *Pvalb* (green) on a brain tissue section (S1BF; 20 μm thick) does not indicate substantial overlap between these markers. However, as shown in (**C**), the use of CCK-Cre with a tdTomato reporter (in this case, CCK-Cre; Dlx5/6-Flpe; Ai65) in combination with IHC for PV (green) results in substantial labeling of PV INs (white arrows). (**D**) CCK-Cre cumulative recombination can also label putative NGFC (NPY+), as shown in an intersectional CCK-Cre; Htr3a-Flpo; Ai65; NPY-hrGFP cross (white arrows indicate tdTomato/NPY+ cells). Scale bars indicate 100 μm.

#### Fig. S7: Comparison of 5HT3aR lines in S1BF

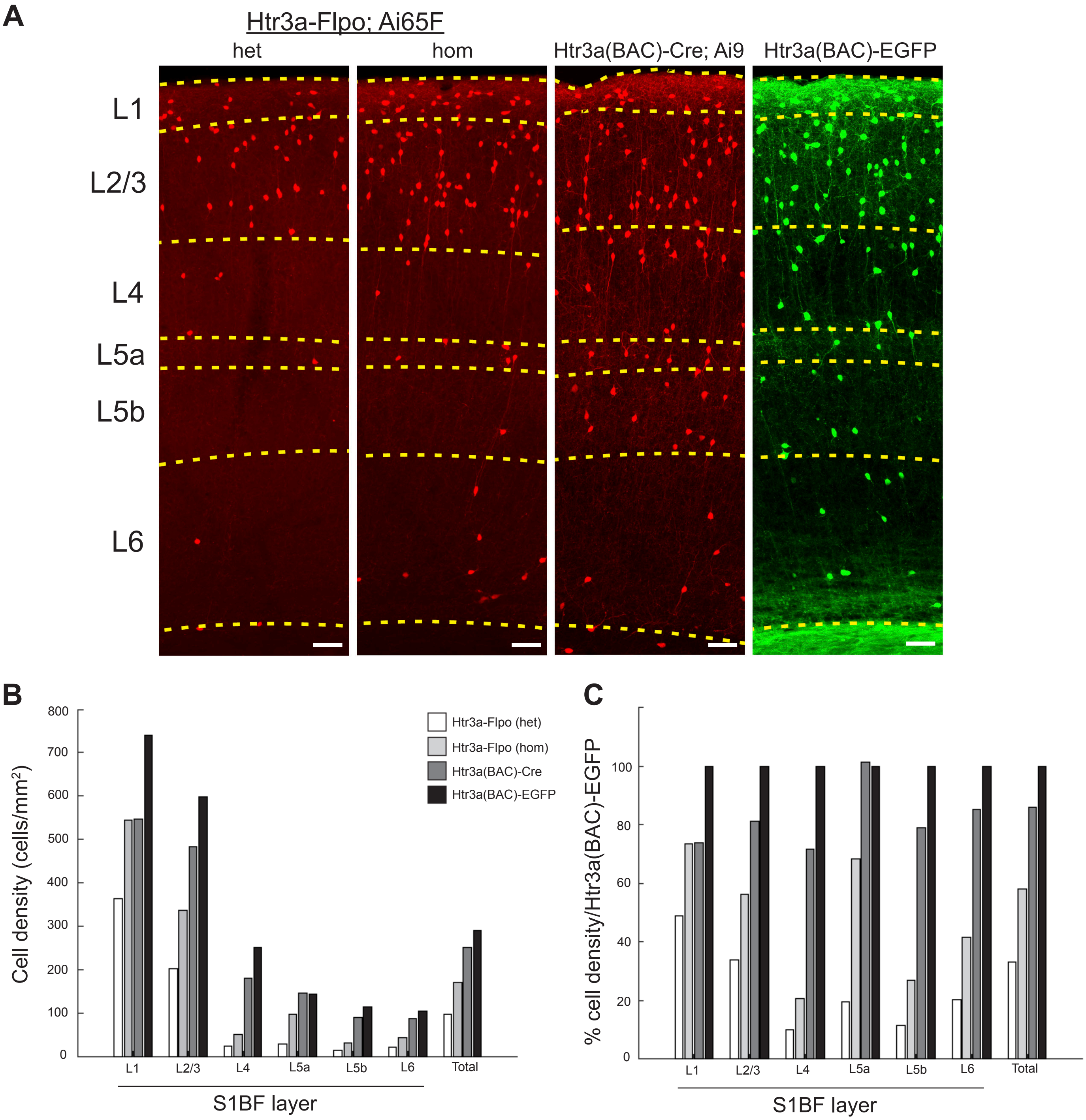

**Figure S7: Comparison of 5HT3aR lines in S1BF.** (**A**) Images of cell labeling in S1BF obtained with one copy of Htr3a-Flpo (het: Htr3a-Flpo/+; Ai65F/+) or two copies (hom: Htr3a-Flpo/Htr3a-Flpo; Ai65F/+), compared with Htr3a(BAC)-Cre; Ai9 and Htr3a(BAC)-EGFP. Dashed lines indicate approximate layer boundaries. (**B**) The density of cell labeling (cells/mm$^2$) obtained from the four Htr3a transgenic labeling strategies across all layers in S1BF. (**C**) The comparative density of cell labeling obtained from the four Htr3a transgenic labeling strategies across all layers in S1BF, normalized to Htr3a(BAC)-EGFP (100%). Overall, one copy of Htr3a-Flpo yielded about 30% of the cell labeling as that seen in Htr3a(BAC)-EGFP animals, with two copies of Htr3a-Flpo doubling this labeling. This observation is consistent with a broad but transient expression of Htr3a mRNA in most CGE INs during embryonic development. The ephemeral nature of Htr3a expression in many CGE INs limits the efficiency of labeling with Htr3a-Flpo, although this can be partially compensated for with two copies of the driver. The Htr3a(BAC)-Cre driver in combination with Ai9 labels ~80% of the cells of the Htr3a(BAC)-EGFP line. Scale bars indicate 100 $\mu$m.
